# Predicting the bacterial host range of plasmid genomes using the language model-based one-class SVM algorithm

**DOI:** 10.1101/2024.08.27.609848

**Authors:** Tao Feng, Xirao Chen, Shufang Wu, Waijiao Tang, Hongwei Zhou, Zhencheng Fang

**Author notes:** To whom correspondence should be addressed., Institutional addresses: 253 Gongye Middle Avenue, Haizhu District, Guangzhou, Guangdong, China, 510280., Hongwei Zhou, Zhencheng Fang. Email addresses: Tao Feng, Xirao Chen, Shufang Wu, Waijiao Tang.

## Abstract

The prediction of the plasmid host range is crucial for investigating the dissemination of plasmids and the transfer of resistance and virulence genes mediated by plasmids. Several machine learning-based tools have been developed to predict plasmid host ranges. These tools have been trained and tested based on the bacterial host records of plasmids in related databases. Typically, a plasmid genome in databases such as NCBI is annotated with only one or a few bacterial hosts, which does not encompass all possible hosts. Consequently, existing methods may significantly underestimate the host ranges of mobilizable plasmids. In this work, we propose a novel method named HRPredict, which employs a word vector model to digitally represent the encoded proteins on plasmid genomes. Since it is difficult to confirm which host a particular plasmid definitely cannot enter, we develop a machine learning approach for predicting whether a plasmid can enter a specific bacterium as a no negative samples learning task. Using multiple one-class SVMs that do not require negative samples for training, the HRPredict predicts the host range of plasmids across 45 families, 56 genera, and 56 species. In the benchmark test set, we constructed reliable negative samples for each host taxonomic unit via two indirect methods, and we found that the *AUC, F1-score, recall, precision*, and *accuracy* of most taxonomic unit prediction models exceeded 0.9. Among the 13 broad-host-range plasmid types, HRPredict demonstrated greater coverage than HOTSPOT and PlasmidHostFinder, thus successfully predicting the majority of hosts previously reported. Through the feature importance calculation for each SVM model, we found that genes closely related to the plasmid host range are involved in functions such as bacterial adaptability, pathogenicity, and survival. These findings provide significant insight into the mechanisms through which bacteria adjust to diverse environments through plasmids.

**Impact Statement:** Plasmids are important vectors for horizontal gene transfer and play a crucial role in regulating bacterial host adaptation to the environment. The spread of plasmid-mediated antibiotic resistance genes and virulence factors is one of the most important public health issues today. Owing to the lack of highly efficient methods for predicting the host range of newly discovered plasmids, especially broad-host-range plasmids, it is difficult to fully elucidate the regulatory role of plasmids in microbial communities and to predict the risk of antibiotic resistance transmission in clinical settings. Existing prediction tools tend to underestimate the host range of mobilizable plasmids. The current paper aims to overcome this limitation. Based on the concept of a “no negative samples learning task,” we propose a new plasmid host range prediction method (i.e., HRPredict) that uses an SVM algorithm based on language models. HRPredict may be a powerful tool that will improve biologists’ understanding of horizontal plasmid transfer and help predict the occurrence and development of bacterial resistance.

**Data Summary:** HRPredict is freely available via https://github.com/FengTaoSMU/HRPredict.

## Introduction

Plasmids are genetic materials that exist outside the bacterial chromosome and can help bacterial hosts acquire environmental adaptability factors [1]. Based on their mobility, plasmids can be categorized into non-mobilizable plasmids and mobilizable plasmids [2-4]. Mobilizable plasmids, especially broad-host-range plasmids, can carry resistance genes, virulence factors, and heavy metal resistance proteins, thereby facilitating their spread among different hosts and increasing the environmental adaptability of their bacterial hosts [5]. Therefore, annotating the host range of plasmids is crucial for studying the dissemination of plasmids and the mechanism of horizontal gene transfer.

Currently, methods for determining the host range of plasmids include experimental and machine learning-based computational approaches. The experimental method of Hi-C sequencing primarily involves crosslinking plasmids with bacterial DNA that are spatially proximate within bacterial cells and obtaining host information for the plasmids through sequencing [6]. However, this method can only identify the current host of some plasmids and is relatively costly. Computational methods include HOTSPOT [7] and PlasmidHostFinder [8]. HOTSPOT, which was developed based on transformers, can predict the hosts of plasmids from the phylum to the species level. PlasmidHostFinder, which was developed based on *k*-mer frequencies and the random forest algorithm, can annotate the bacterial hosts of plasmids from the class level to the species level. These tools have been primarily trained and tested based on the bacterial host records of plasmids in related databases. Generally, for a specific plasmid genome, the database does not record all the possible hosts that the plasmid can enter. For example, when a plasmid genome file is downloaded from NCBI, the file typically records only the taxonomic unit of the bacterial strain from which the plasmid was isolated and sequenced. Therefore, using this information to train and test tools may lead to a significant underestimation of the host range of many plasmids, especially broad-host-range plasmids. Moreover, owing to the high diversity of plasmids, many newly discovered plasmids lack homology with plasmid sequences in existing databases [9,10]. Additionally, the functions of many genes on plasmids remain unclear [10], and our understanding of the molecular mechanisms of plasmid transfer is limited. These factors present significant challenges for mathematical modeling of plasmid sequences during algorithm development.

In recent years, artificial intelligence language models have advanced by treating short amino acid or nucleotide sequences as “words” and treating protein or DNA sequences as sentences [11,12]. Through unsupervised pretraining on large datasets, these models can generate context-based vector representations for each “word”. Compared with traditional mathematical representation methods for biological sequences, this approach can capture deeper abstract information underlying sequences from one-dimensional data and uncover unknown information about new sequences. Currently, language models are widely used in various biological sequence prediction methods, such as protein three-dimensional structure prediction [13], gene function annotation [14,15], low-homology protein searches [16], plasmid typing [17], and probiotic prediction [18]. By overcoming the challenges posed by low sequence homology and unknown sequence functions, language models also present opportunities for annotating the host range of plasmids.

In this work, we introduce a tool named HRPredict, which is used to predict bacterial hosts for plasmids across 45 families, 56 genera, and 56 species. HRPredict accepts fasta files of complete plasmid genomes as input and outputs the host range at the family, genus, and species levels. By utilizing the language model-based word vectors of ProtVec [12], HRPredict generates word vector distance-based representations of plasmid-encoded proteins. Considering that we can only obtain information from databases about which bacteria a plasmid can enter but no information about which bacteria a plasmid absolutely cannot enter, we employed a one-class SVM algorithm, which does not require negative samples, to train a classifier for each taxonomic unit, thereby increasing the tool’s sensitivity. On the other hand, to assess whether the tool produces false positive predictions, we need to test the tool using negative samples. Based on biological principles, we constructed reliable negative samples for the test set via two indirect methods to evaluate the tool performance. Testing showed that HRPredict performs well and is better at predicting the known hosts of broad-host-range plasmids.

## METHODS

### The workflow for predicting plasmid host ranges using HRPredict

As shown in Figure 1A, HRPredict takes fasta files of complete plasmid genomes as input. For each plasmid genome sequence, HRPredict annotates its encoded proteins via Prokka [19]. Subsequently, the tool generates a protein feature vector 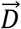 for a plasmid genome based on ProtVec [12], a pretrained protein language model. Through a series of prediction modules trained with one-class SVMs, HRPredict outputs the host range of the input plasmids at the family, genus, and species levels.

**Figure 1.**
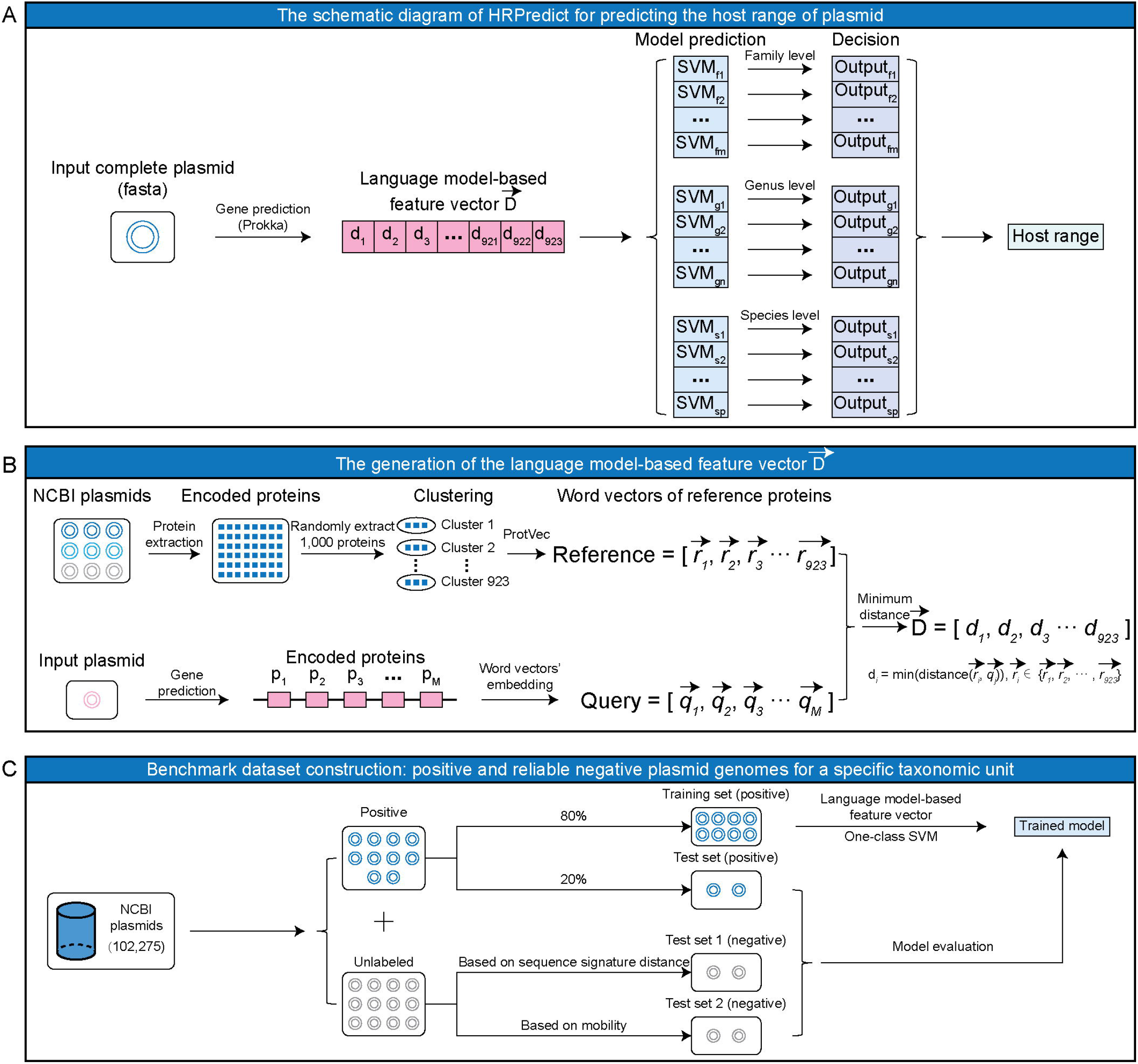
Development workflow of HRPredict. (A) Diagram illustrating the process of using HRPredict to predict plasmid host ranges. (B) The process of generating plasmid genome feature vectors based on a language model. (C) Generation of benchmark datasets and model training for a specific host taxonomic unit.

### Generation of the language model-based feature vector 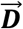

We downloaded complete plasmid genomes from NCBI and excluded some known broad-host-range plasmids that serve as independent datasets for subsequent tool application. This resulted in a total of 102,544 plasmid genomes (Table S1). For each plasmid genome, we generated its feature representation in accordance with the following process (Figure 1B):

a. Generation of the reference protein feature vector set. All encoded protein sequences were extracted from plasmid genomes downloaded from NCBI. We randomly selected *n* (*n*=1,000) protein sequences (the choice of *n* will be described in the Discussion section) and used CD-HIT [20] to cluster these proteins with 90% similarity, thus yielding 923 non-redundant protein sequences as reference plasmid proteins. For each reference protein sequence, we use the pretrained ProtVec model to map every adjacent three amino acids on a protein to a 100-dimensional word vector. The average vector of all three amino acid word vectors in the sequence is then used as the feature vector for that protein. This results in a reference feature vector set for 923 non-redundant protein sequences, namely, 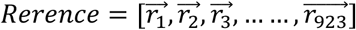 (Figure 1B).
b. Generation of the input plasmid protein feature vector. For each plasmid genome, Prokka is used to annotate its encoded protein sequences, and ProtVec is employed to calculate the vector representation of each protein sequence as mentioned above. This process generates a feature vector set for all protein sequences encoded by the input plasmid, namely, 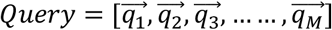 (Figure 1B), where *M* denotes the number of proteins on the plasmid genome.
c. Generation of the feature vector for the input plasmid. As shown in Figure 1B, each input plasmid is eventually represented by a 923-dimensional vector 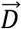. The *i*th element of vector 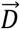 represents the shortest Euclidean distance between all vectors in the Query set and the *i*th vector in the Reference set. Thus, the vector 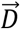 provides a spatial representation of the plasmid-encoded genes based on the distances between word vectors.

### Positive samples of the training and test sets for a specific host taxonomic unit

The host taxonomic information of each plasmid genome was extracted based on the annotations from its GenBank file. We selected taxonomic units with more than 100 plasmids at the family, genus, and species levels for subsequent analysis. Uncertain taxa, such as those labeled as unclassified, were not included in the model. This resulted in a total of 45 families, 56 genera, and 56 species, including many clinically significant drug-resistant bacteria. For each plasmid genome, the feature vector 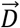 was generated, and plasmids with the same feature vectors were removed. Subsequently, 80% of the plasmid genomes within this host taxonomic unit were randomly selected to generate positive samples for the training set, whereas the remaining 20% were used as positive samples for the test set (Figure 1C).

### Two strategies for selecting reliable negative samples for specific taxonomic units in the test set

a. Based on the sequence signature distance. Owing to genome amelioration, the sequence characteristics of plasmid DNA, such as GC content and codon usage preference, tend to resemble those of the host bacterial chromosomal DNA [21]. Conversely, if a plasmid has never been hosted by a particular bacterium, its sequence characteristics will significantly differ from those of that bacterium. Based on Suzuki’s approach [21], we used DNA 4-mer frequency vectors to measure the sequence similarity between different plasmids. For a given taxon A, we first calculate the average 4-mer frequency of plasmids from taxon A, and then compute the Euclidean distance between the 4-mer frequency of each plasmid not labeled as belonging to A and the average frequency vector of A plasmids. We then selected the plasmids from other taxon with the largest Euclidean distances, matching the number of plasmids from taxon A, to serve as the negative samples for taxon A.
b. Based on mobility. For taxon A, non-mobilizable plasmids from other taxonomic units can serve as negative samples. The presence of relaxase is a key marker that differentiates mobilizable plasmids from non-mobilizable ones. Using our previous method [17], we identified non-mobilizable plasmids by searching for relaxase in the plasmid genomes. For a given taxon A, we randomly selected non-mobilizable plasmids from other taxa, matching the number of plasmids in taxon A, to use as negative samples for taxon A.

### Module Training and Integration

For each specific host taxonomic unit, we used the *svm* function (parameters: type = “one-classification”, nu = 0.01, scale = TRUE, kernel = “radial”) from the R package *e1071* to train a one-class SVM without negative samples. In the final version of HRPredict, we linked these modules in a sequence following the order of family, genus, and species. Each input plasmid first undergoes evaluation by the higher-level classifier. If a plasmid is identified as belonging to a particular taxonomic unit, it is then passed to the next-level classifier specific to that taxonomic unit.

## RESULTS

### Performance evaluation of HRPredict

We evaluate the performance of the discrimination modules of each taxonomic unit in HRPredict using the benchmark test set. Since the negative samples in the test set are generated by two different methods, we separately observe the performance of each module under the two types of test data (Figure 2). We first evaluate the *AUC* of different modules and identify the threshold that maximizes the Youden index for each module using the ROC curve as the default judgment threshold. At this threshold, we further evaluate other metrics, including *accuracy*=(TP+TN)/(TP+TN+FN+FP), *recall*=TP/(TP+FN), *precision*=TP/(TP+FP), and *F1-score*=2×*recall*×*precision*/(*recall*+*precision*). Here, TP, TN, FN, and FP represent the number of true positive, true negative, false negative, and false positive predictions, respectively. We found that, in most cases, the different modules of HRPredict achieved good performance, with values greater than 0.9 for the majority of evaluation metrics. Moreover, when the modules were tested on the test set generated based on mobility information for negative samples, the values of various metrics were lower than those on the test set based on sequence signature information, especially the precision metric, suggesting that there may be some false positives in the predictions. We primarily determine whether a plasmid is mobilizable by the presence of relaxase on the plasmid. Although this criterion is widely accepted by biologists, some relaxases may be missed because of the high diversity of relaxase sequences. In fact, many new relaxases have been discovered in recent years [22,23], and the sequences of these new relaxases differ from those of existing relaxases. Inadequate annotation of relaxases may lead to mobilizable plasmids being annotated as non-mobilizable. Therefore, the false positive predictions observed in the performance evaluation may not necessarily be incorrect and could originate from overlooked mobilizable plasmids. In the released version of HRPredict, we used the threshold confirmed by sequence signature information as the final decision threshold. The detailed performance metrics for different taxon modules are shown in Figures S1-S3. The results indicate that the performance of the modules for some common resistant bacteria, such as *Klebsiella pneumoniae* and *Escherichia coli*, is relatively low. This finding implies that these resistant bacteria might have more complex plasmid exchange mechanisms, thus making prediction more challenging. Nevertheless, the modules for these resistant bacteria still achieved a commendable *AUC* greater than 75%.

**Figure 2.**
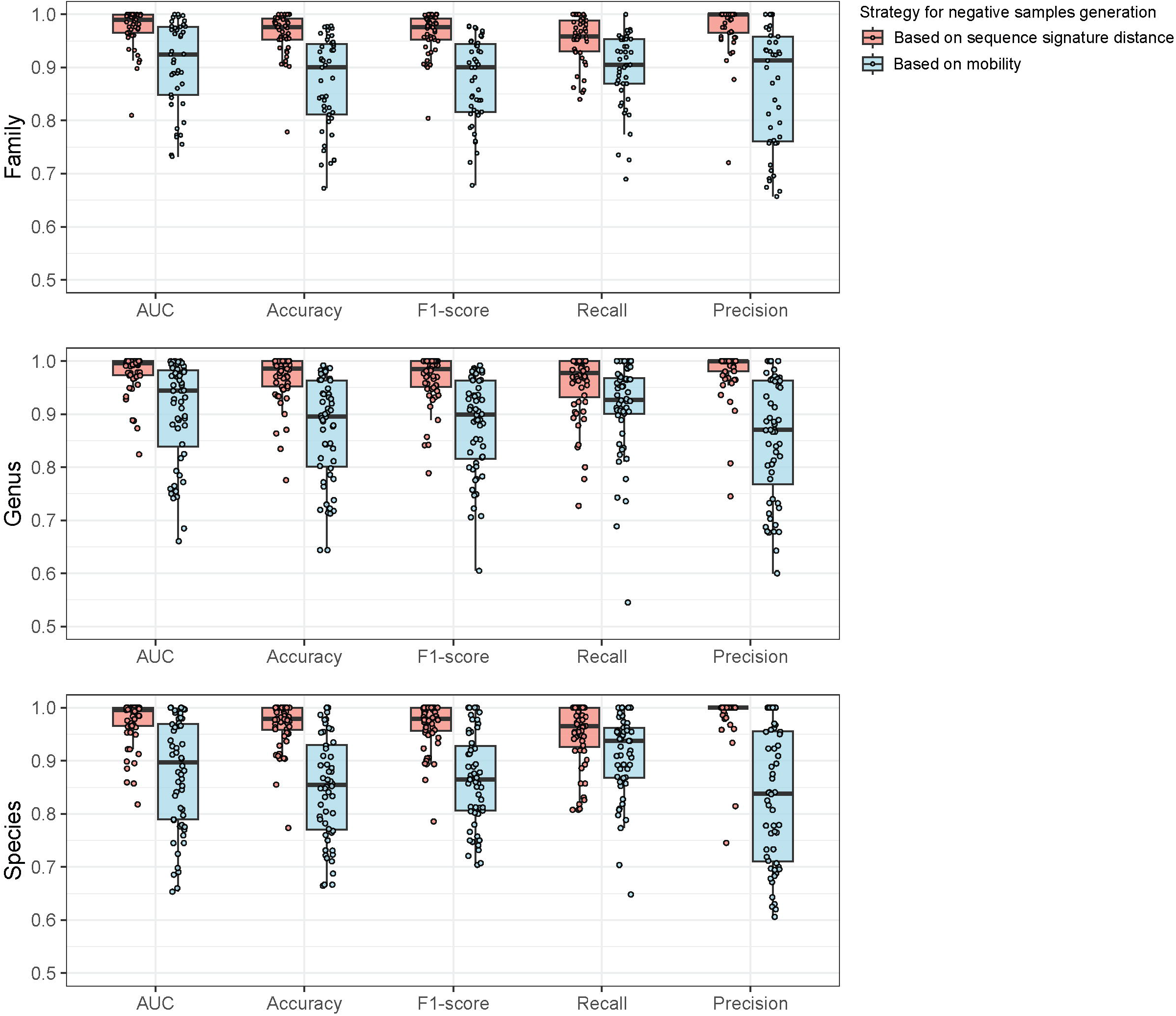
Performance evaluation of one-class SVM modules at different taxonomic unit levels in HRPredict. Each point in the box plot represents a one-class SVM for a specific taxonomic unit. The performance of each module is observed separately under the two types of test data.

### Application and comparison of broad-host-range plasmids

From the broad-host-range plasmids we collected, we selected plasmid types with at least three reported host genera, as shown in Table S2. This selection was used to evaluate the performance of HRPredict (released version), HOSTPOT, and PlasmidHostFinder in predicting the host range of broad-host-range plasmids at the genus level. For each plasmid, we collected multiple genomes, as shown in Table S2. In our evaluation, if a tool predicted that any genome of a particular plasmid type could infect a specific genus, we considered that the tool had successfully identified that genus as a host for that plasmid type. During the evaluation process, we excluded plasmid genomes that were artificially synthesized or modified.

We found that HRPredict has the highest coverage, averaging coverage of about 70% of the reported hosts. In contrast, HOSTPOT and PlasmidHostFinder could not predict most of the hosts of these plasmids, with an average coverage of less than 25% (Figure 3A). This finding further demonstrates the advantage of the HRPredict tool in terms of predicting the range of plasmid hosts. Meanwhile, we noticed that each tool predicted some hosts that had not been found reported as far as we known. Without additional evidence, determining the accuracy of these extra predictions is challenging. Nevertheless, we quantitatively compared the predictions of reported hosts and the additional host predictions among the different tools. We defined a ratio called CA, which is the ratio of the coverage of reported hosts for a certain type of plasmid by the tool to the proportion of additional hosts predicted by the tool. The ratio is calculated as follows: 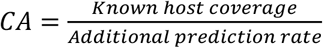. We found that the average CA value did not vary significantly across different tools, and HRPredict showed relatively stable values across different types of plasmids (Figure 3B). This finding suggests that many broad-host-range plasmids may have additional bacterial hosts that have not been identified, and our tool could provide some clues for identifying these new hosts.

**Figure 3.**
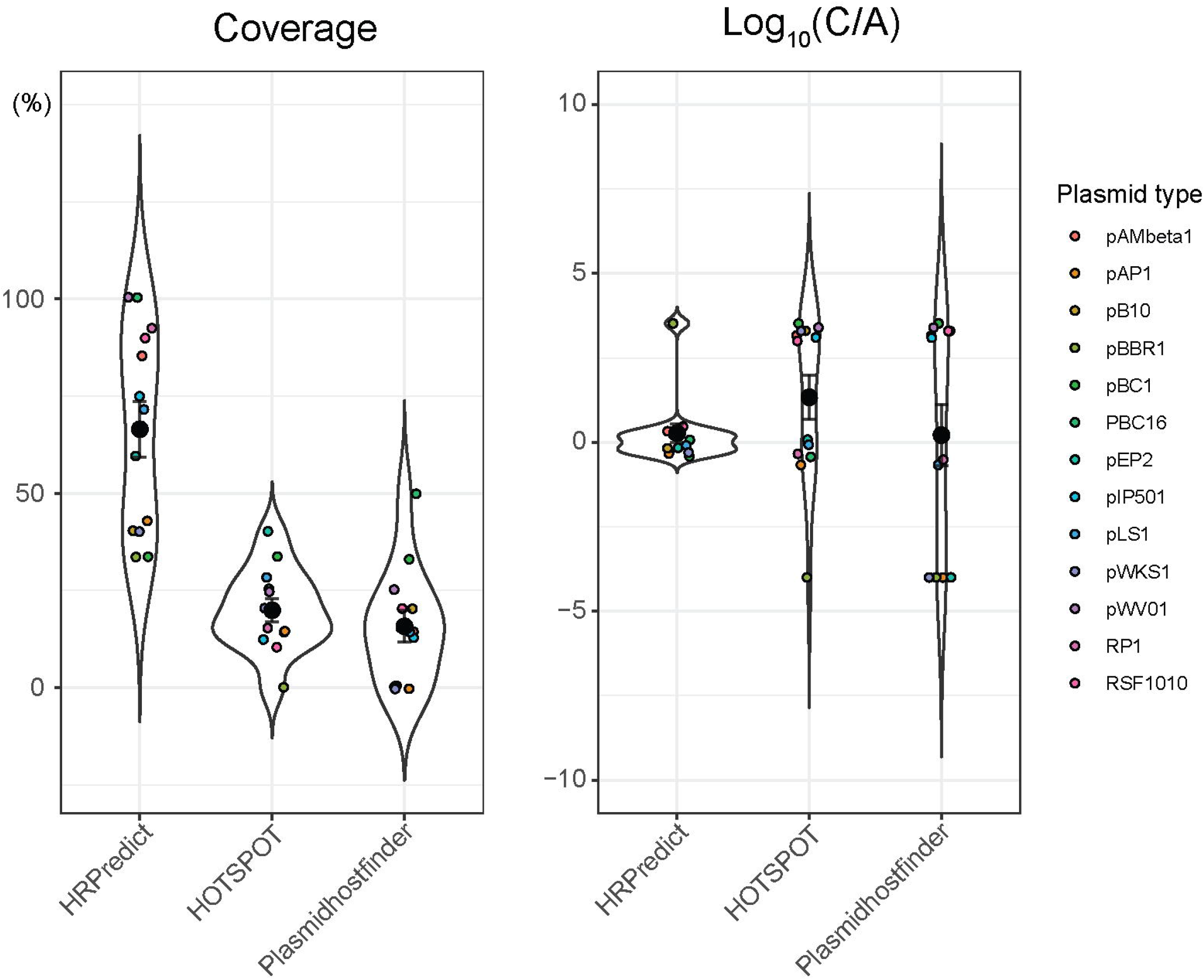
Application and comparison of broad-host-range plasmids with known host ranges. (A) The prediction coverage rates of reported hosts (at the genus level) for different types of broad-host-range plasmids by each tool. (B) CA values for the predicted host ranges of different plasmid types according to each tool. For clarity, we have adjusted the axis to 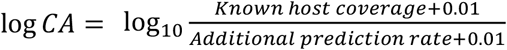. Each small dot in the figure represents the value for a specific type of plasmid, and the larger dots indicate the average values.

### Feature importance of each classifier at different taxonomic levels

Among the 923-dimensional features input into each module, each dimension represents a specific reference protein. For each taxonomic level’s SVMs, we calculated the permutation importance for each feature. The top 10 most important features for each SVM were then collected based on these absolute values. From these features, we identified the 15 most frequently occurring features as the most important features at that level, as shown in Figure 4.

**Figure 4.**
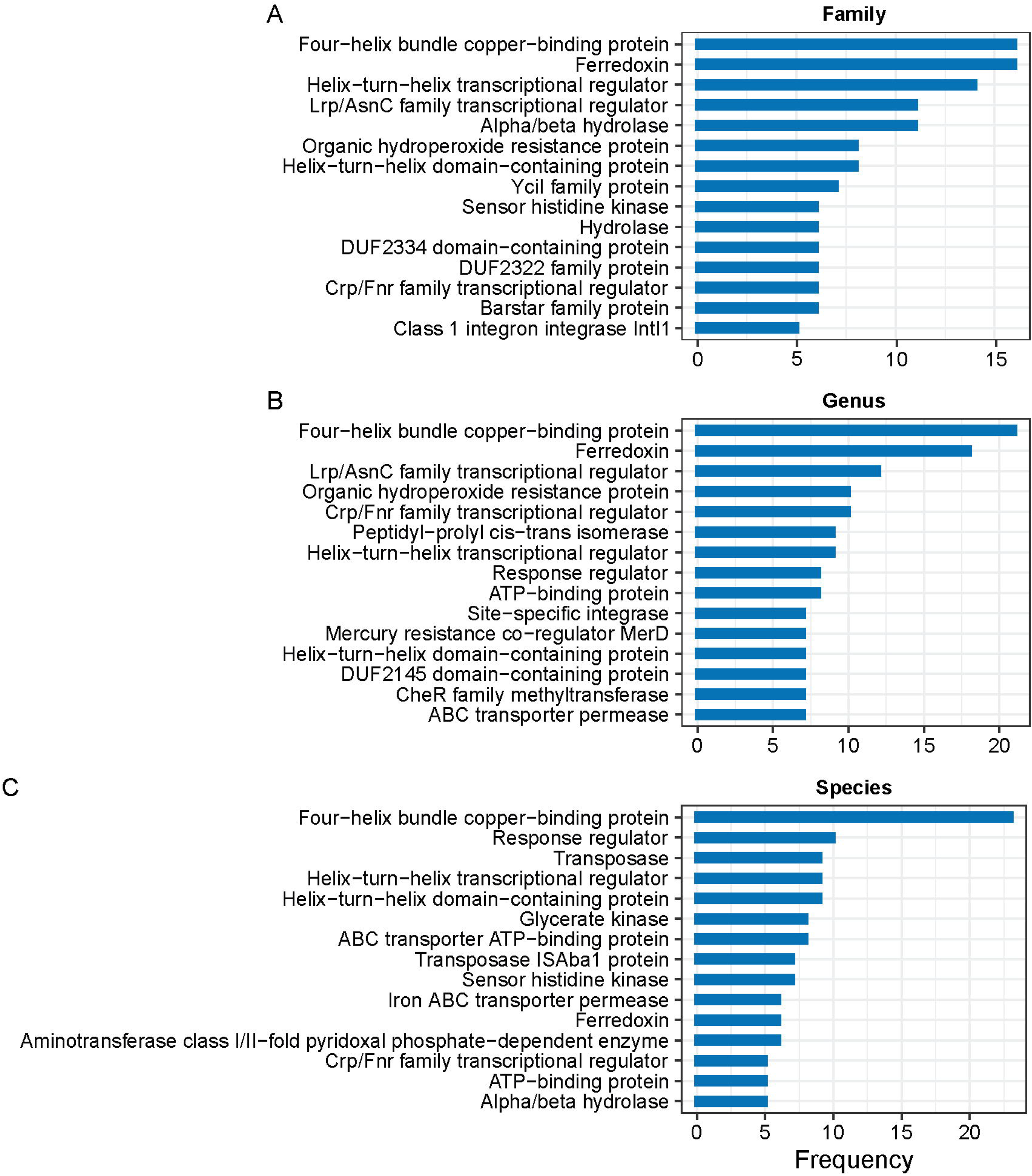
The top 15 most important features for each taxonomic level.

We found that the most important features identified above collectively indicate adaptive mechanisms that bacteria employ to thrive in diverse environments, enhance their survival, and contribute to their pathogenicity through plasmids. For example, the four-helix bundle copper-binding protein and ferredoxin are the two genes most strongly associated with the host range. These proteins are involved in metal ion binding and redox reactions, respectively. They play roles in electron transport and maintaining the cellular redox balance, which are important for various metabolic processes and adaptation to different host environments. The response regulator also ranks among the top two at one of the taxonomic levels. The response regulator is a crucial component of bacterial two-component regulatory systems (TCSs), which enable bacteria to sense and respond to environmental changes [24]. Moreover, at the species level, the transposase and transposase ISAba1 proteins are strongly correlated with the host range. These genes play significant roles in the mobility of genetic elements within bacterial genomes, facilitating the spread of antibiotic resistance genes and virulence factors and thus enhancing bacterial pathogenicity. Specifically, the ISAba1 transposase is frequently found in multidrug-resistant *Acinetobacter baumannii* strains, increasing their pathogenicity and resistance to antibiotics [25]. Although it is challenging to determine whether the presence of these genes has a direct causal relationship with the host range of plasmids, these findings can aid in further understanding the regulatory mechanisms and functions of plasmids in bacteria.

## Discussion

Herein, we employed a one-class SVM without negative samples to train the classifiers. This approach better reflects real-world scenarios since databases only record the known hosts of a plasmid without indicating bacteria that the plasmid cannot enter. In the application cases of broad-host-range plasmids, this method has also demonstrated its ability to help the algorithm predict more potential hosts. However, owing to the lack of negative sample constraints, the tool is prone to false positives. We used two indirect methods to construct negative samples in the test set and ultimately selected the decision threshold for the tool based on negative samples derived from sequence signature information. Importantly, sequence signatures are based on nucleic acid information, reflecting characteristics such as GC content and codon usage bias. In contrast, the feature vectors used in HRPredict are based on protein information measured by a language model, and these two types of information are relatively independent. Therefore, the use of such negative sample test data can objectively reflect the tool’s performance.

Using all plasmid proteins from NCBI to construct the reference set would result in excessively high feature vector dimensions, leading to overfitting and increased training difficulty. To address this, we employed random sampling and clustering of plasmid proteins to reduce the dimensionality of the feature vectors. We sampled the plasmid proteins in varying quantities and then clustered them based on 90% similarity to extract the reference proteins. Using these reference proteins, we constructed the feature vectors and repeatedly trained the modules of HRPredict. We found that when the number of sampled proteins reached 1,000, the *AUC, F1-score, recall, precision*, and *accuracy* of the modules stabilized (Figure S4). Therefore, we set the number of sampled proteins to 1,000.

HRPredict can determine the host range of plasmids based solely on the information of encoded proteins. However, in real-world scenarios, whether a plasmid can enter a specific bacterium is influenced not only by the encoded genes but also by numerous external factors, such as selective pressure. Therefore, we believe that HRPredict’s predictions should be understood as identifying all potential hosts that a plasmid can enter given its encoded gene functions but not necessarily the bacteria it will enter in a specific environment. Predicting the transfer pathways of antibiotic-resistant plasmids under antibiotic stimulation is crucial for monitoring clinical antibiotic-resistant bacteria. In future work, we plan to collect plasmid transfer data under specific conditions, such as antibiotic intervention, incorporate environmental factors into our model, and use multimodal information to make more precise predictions of plasmid horizontal transfer. Moreover, the range of hosts that HRPredict can predict does not encompass all taxa. We selected the corresponding taxonomic unit for analysis based on the amount of plasmid genome data available. Although the taxonomic unit we included covers most of the important bacteria, in our future work, we will attempt to use techniques designed for small sample sizes, such as transfer learning, to enable HRPredict to make predictions for a broader range of taxa.

---

**Figure S1**. The performance of discriminators for each taxonomic unit at the family level.

**Figure S2**. The performance of discriminators for each taxonomic unit at the genus level.

**Figure S3**. The performance of discriminators for each taxonomic unit at the species level.

**Figure S4**. The performance of each module under different quantities of random plasmid protein sampling. The generation of negative samples in these analyses is based on sequence signature distance.

**Table S1**. The plasmid accessions used in the construction of HRPredict.

**Table S2**. The plasmid types and genome accessions used to compare the performance of various tools in predicting broad-host-range plasmids are listed in the table. The reported host genera and their corresponding host ranges are also included.

## References

1. Rodríguez-Beltrán J, DelaFuente J, León-Sampedro R, MacLean RC, San Millán Á. Beyond horizontal gene transfer: the role of plasmids in bacterial evolution. Nat Rev Microbiol. 2021, 19(6):347–359.

2. Francia MV, Varsaki A, Garcillán-Barcia MP, Latorre A, Drainas C, et al. A classification scheme for mobilization regions of bacterial plasmids. FEMS Microbiol Rev. 2004, 28(1):79–100

3. Smillie C, Garcillán-Barcia MP, Francia MV, Rocha EP, de la Cruz F. Mobility of plasmids. Microbiol Mol Biol Rev. 2010, 74(3):434–52.

4. Garcillán-Barcia MP, Alvarado A, de la Cruz F. Identification of bacterial plasmids based on mobility and plasmid population biology. FEMS Microbiol Rev. 2011, 35(5):936–56.

5. Arnold BJ, Huang IT, Hanage WP. Horizontal gene transfer and adaptive evolution in bacteria. Nat Rev Microbiol. 2022, 20(4):206–218.

6. Brito IL. Examining horizontal gene transfer in microbial communities. Nat Rev Microbiol. 2021, 19(7):442–453.

7. Ji Y, Shang J, Tang X, Sun Y. HOTSPOT: hierarchical host prediction for assembled plasmid contigs with transformer. Bioinformatics. 2023 May, 39(5):btad283.

8. Aytan-Aktug D, Clausen PTLC, Szarvas J, Munk P, Otani S, et al. PlasmidHostFinder: Prediction of Plasmid Hosts Using Random Forest. mSystems. 2022, 7(2):e0118021.

9. Fang Z, Zhou H. Identification of the conjugative and mobilizable plasmid fragments in the plasmidome using sequence signatures. Microb Genom. 2020, 6(11):mgen000459.

10. Fang Z, Tan J, Wu S, Li M, Wang C, Liu Y, Zhu H. PlasGUN: gene prediction in plasmid metagenomic short reads using deep learning. Bioinformatics. 2020, 36(10):3239–3241.

11. Ng, P. dna2vec: Consistent vector representations of variable-length k-mers. 2017; arXiv preprint arXiv:1701.06279.

12. Asgari E, Mofrad MR. Continuous Distributed Representation of Biological Sequences for Deep Proteomics and Genomics. PLoS One. 2015, 10(11):e0141287.

13. Abramson J, Adler J, Dunger J, Evans R, Green T, Pritzel A, Ronneberger O, Willmore L, Ballard AJ, Bambrick J, Bodenstein SW, Evans DA, Hung CC, O’Neill M, Reiman D, Tunyasuvunakool K, Wu Z, Žemgulytė A, Arvaniti E, Beattie C, Bertolli O, Bridgland A, Cherepanov A, Congreve M, Cowen-Rivers AI, Cowie A, Figurnov M, Fuchs FB, Gladman H, Jain R, Khan YA, Low CMR, Perlin K, Potapenko A, Savy P, Singh S, Stecula A, Thillaisundaram A, Tong C, Yakneen S, Zhong ED, Zielinski M, Žídek A, Bapst V, Kohli P, Jaderberg M, Hassabis D, Jumper JM. Accurate structure prediction of biomolecular interactions with AlphaFold 3. Nature. 2024, 630(8016):493–500.

14. Zhu YH, Zhang C, Yu DJ, Zhang Y. Integrating unsupervised language model with triplet neural networks for protein gene ontology prediction. PLoS Comput Biol. 2022, 18(12):e1010793.

15. Wu J, Ouyang J, Qin H, Zhou J, Roberts R, et al. PLM-ARG: antibiotic resistance gene identification using a pretrained protein language model. Bioinformatics. 2023, 39(11):btad690.

16. Liu W, Wang Z, You R, Xie C, Wei H, et al. PLMSearch: Protein language model powers accurate and fast sequence search for remote homology. Nat Commun. 2024, 15(1):2775.

17. Feng T, Wu S, Zhou H, Fang Z. MOBFinder: a tool for mobilization typing of plasmid metagenomic fragments based on a language model. Gigascience. 2024, 13:giae047.

18. Wu S, Feng T, Tang W, Qi C, Gao J, et al. metaProbiotics: a tool for mining probiotic from metagenomic binning data based on a language model. Brief Bioinform. 2024, 25(2):bbae085.

19. Seemann T. Prokka: rapid prokaryotic genome annotation. Bioinformatics. 2014, 30(14):2068–9.

20. Li W, Godzik A. Cd-hit: a fast program for clustering and comparing large sets of protein or nucleotide sequences. Bioinformatics. 2006, 22(13):1658–9.

21. Suzuki H, Yano H, Brown CJ, Top EM. Predicting plasmid promiscuity based on genomic signature. J Bacteriol. 2010, 192(22):6045–55.

22. Wisniewski JA, Traore DA, Bannam TL, Lyras D, Whisstock JC, Rood JI. TcpM: a novel relaxase that mediates transfer of large conjugative plasmids from Clostridium perfringens. Mol Microbiol. 2016, 99(5):884–96.

23. Ramachandran G, Miguel-Arribas A, Abia D, Singh PK, Crespo I, Gago-Córdoba C, Hao JA, Luque-Ortega JR, Alfonso C, Wu LJ, Boer DR, Meijer WJ. Discovery of a new family of relaxases in Firmicutes bacteria. PLoS Genet. 2017, 13(2):e1006586.

24. Rajeev L, Luning EG, Dehal PS, Price MN, Arkin AP, Mukhopadhyay A. Systematic mapping of two component response regulators to gene targets in a model sulfate reducing bacterium. Genome Biol. 2011, 12(10):R99.

25. Turton JF, Ward ME, Woodford N, Kaufmann ME, Pike R, Livermore DM, Pitt TL. The role of ISAba1 in expression of OXA carbapenemase genes in Acinetobacter baumannii. FEMS Microbiol Lett. 2006, 258(1):72–7.

