## Supplementary materials for "Predicting the bacterial host range of plasmid genomes using the language model-based one-class SVM algorithm"

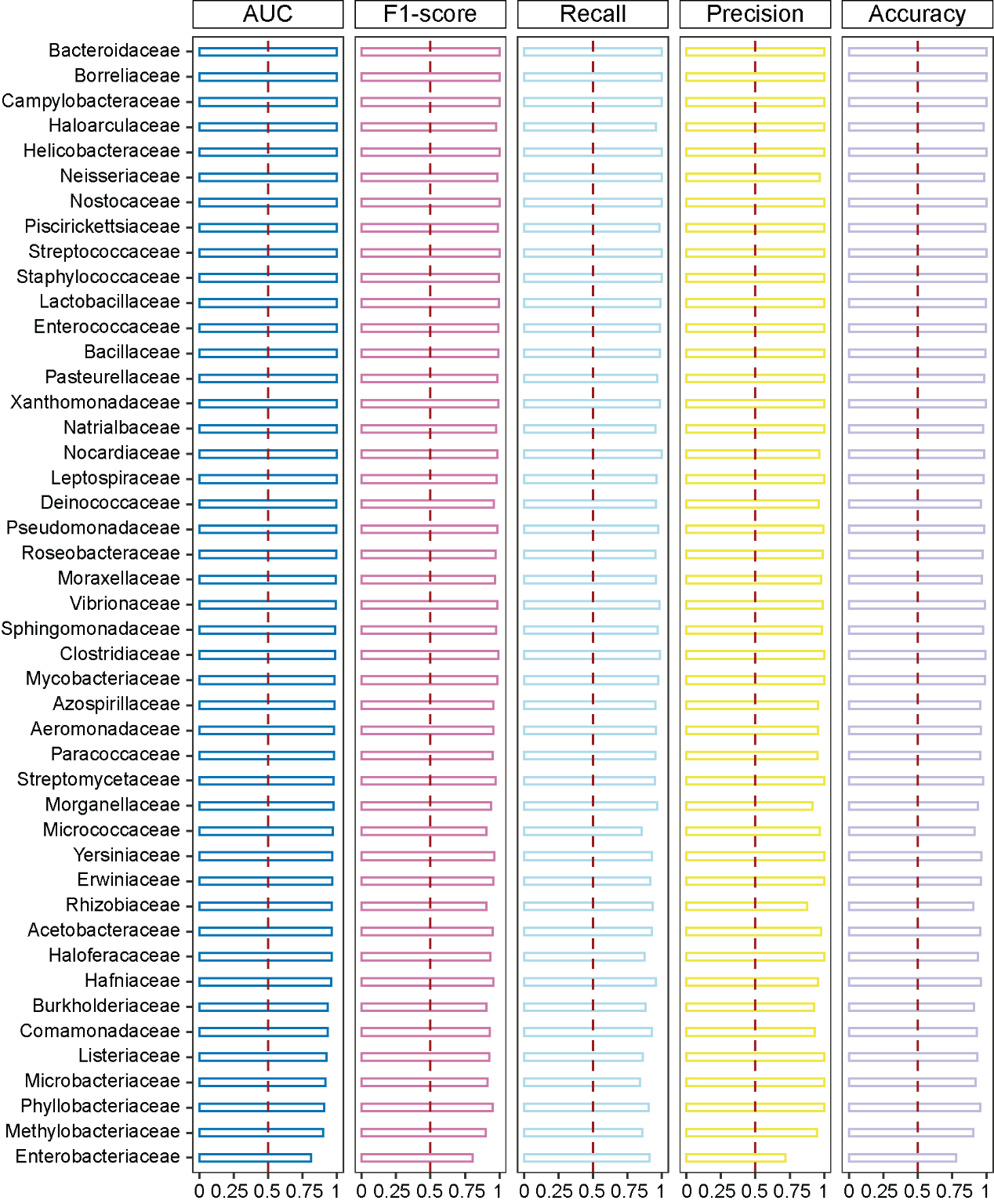

**Figure S1**. The performance of discriminators for each taxonomic unit at the family level.

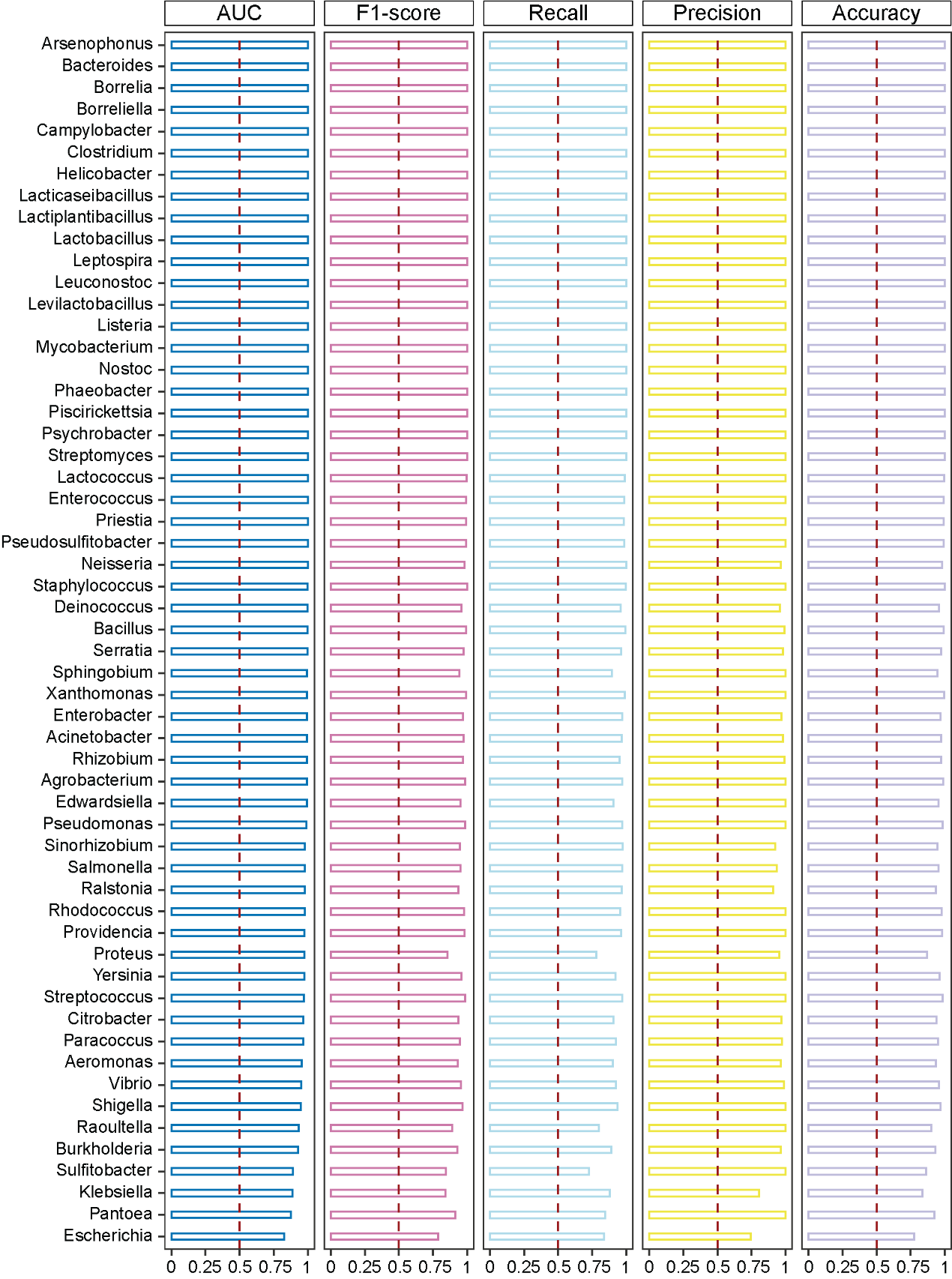

**Figure S2**. The performance of discriminators for each taxonomic unit at the genus level.

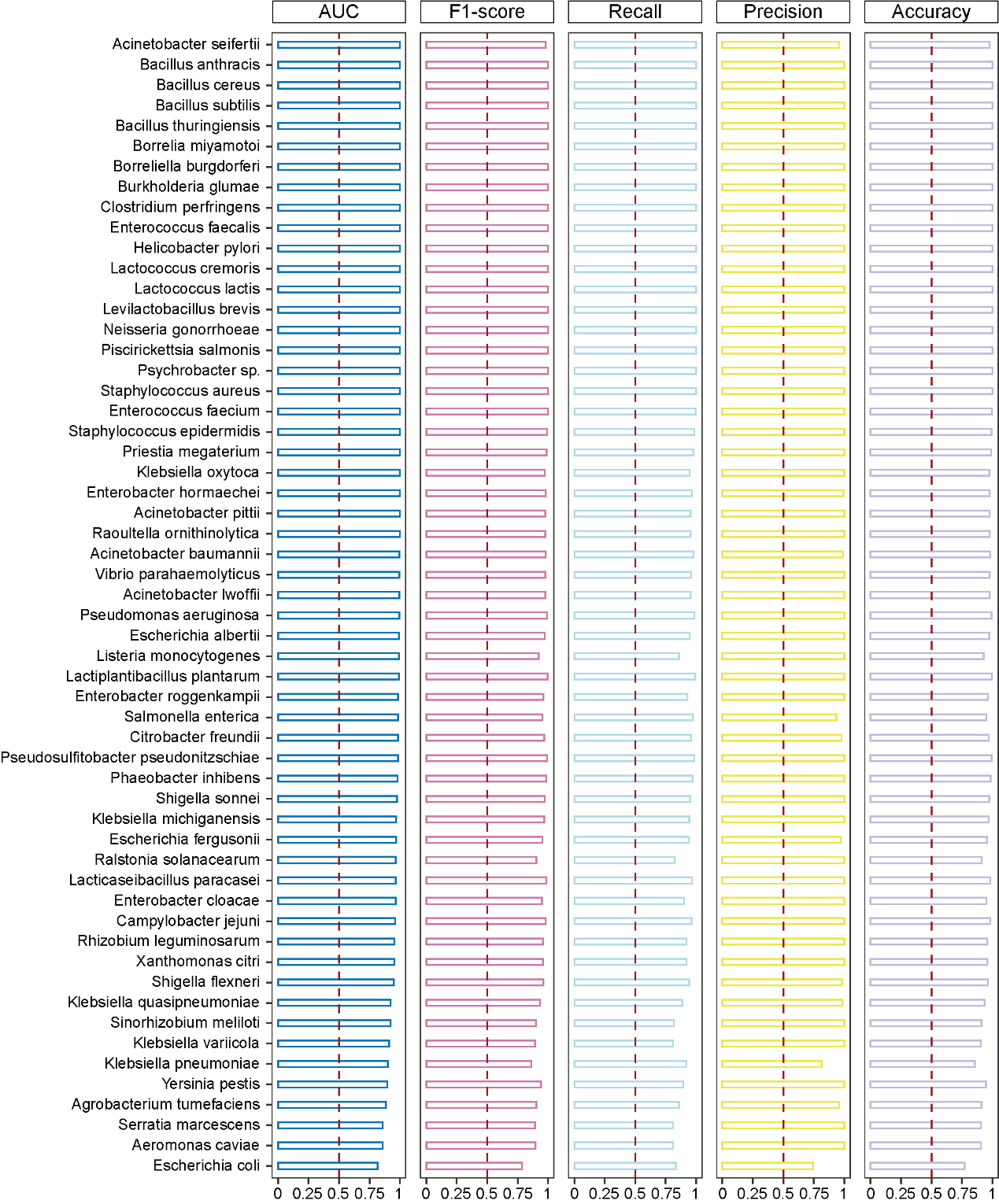

**Figure S3**. The performance of discriminators for each taxonomic unit at the species level.

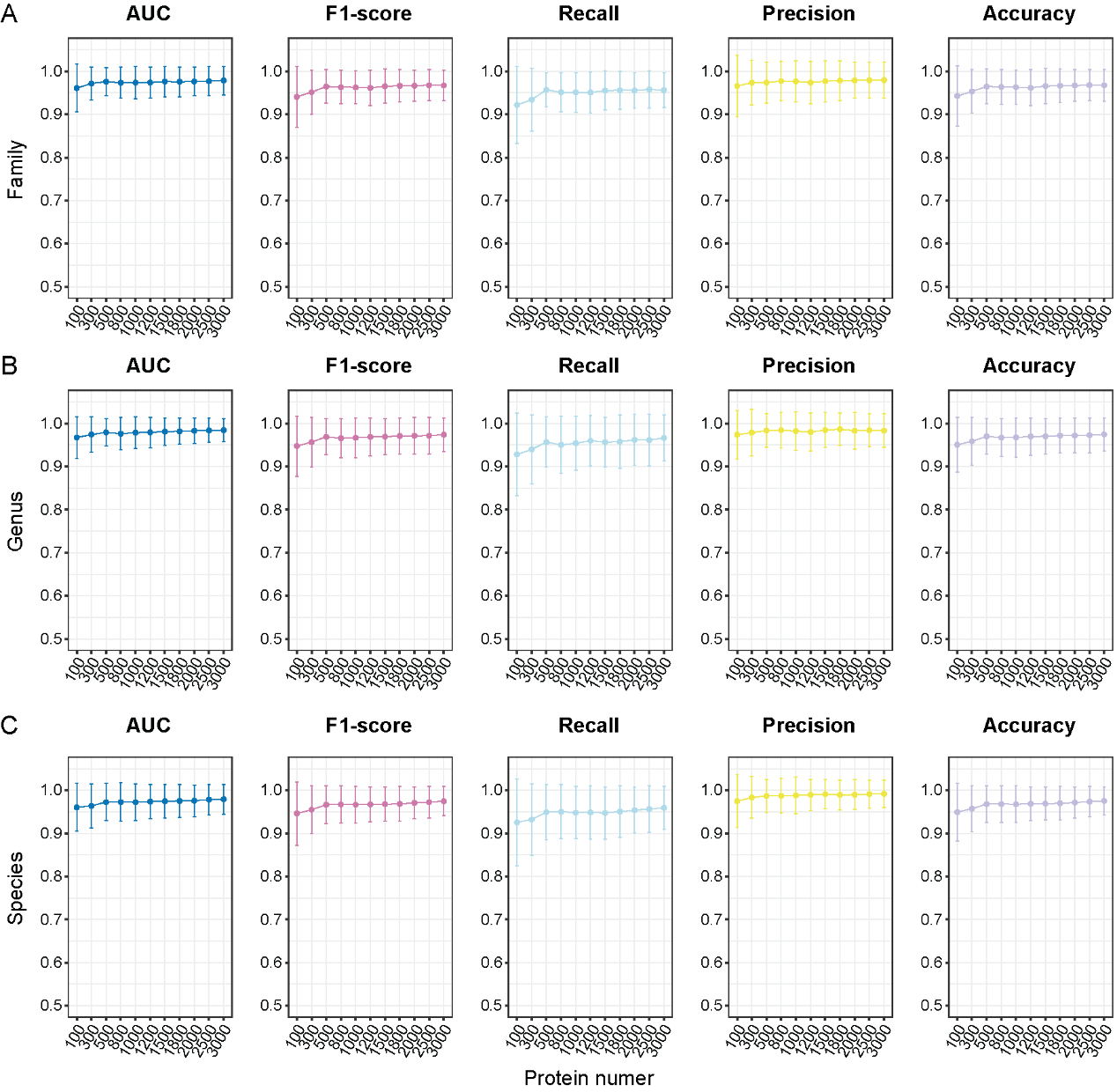

**Figure S4**. The performance of each module under different quantities of random plasmid protein sampling. The generation of negative samples in these analyses is based on plasmid sequence signature distance.
